# Bayesian modeling disentangles language versus executive control disruption in stroke

**DOI:** 10.1101/2023.08.06.552147

**Authors:** Gesa Hartwigsen, Jae-Sung Lim, Hee-Joon Bae, Kyung-Ho Yu, Hugo J. Kuijf, Nick A. Weaver, J. Matthijs Biesbroek, Jakub Kopal, Danilo Bzdok

**Affiliations:** Max Planck Institute for Human Cognitive and Brain Sciences Leipzig, Germany; Wilhelm Wundt Institute for Psychology, Leipzig University, Leipzig, Germany; Department of Neurology, Asan Medical Center, University of Ulsan College of Medicine, Seoul, Republic of Korea; Department of Neurology, Seoul National University Bundang Hospital, Seoul National University College of Medicine, Seongnam, Republic of Korea; Department of Neurology, Hallym University Sacred Heart Hospital, Hallym University College of Medicine, Anyang, Republic of Korea; Image Sciences Institute, UMC, Utrecht, the Netherlands; Department of Neurology, UMC Utrecht Brain Center, University Medical Center Utrecht, Utrecht, The Netherlands; Department of Neurology, Diakonessenhuis Hospital, Utrecht, The Netherlands; Department of Biomedical Engineering, McConnell Brain Imaging Centre, Montreal Neurological Institute, Faculty of Medicine, McGill University, Montreal, Canada; Mila - Quebec Artificial Intelligence Institute, Montreal, Canada

**Keywords:** speech, control, lateralization, domain-general, lesion

## Abstract

Stroke is the leading cause of long-term disability worldwide. Incurred brain damage disrupts cognition, often with persisting deficits in language and executive capacities. Despite their clinical relevance, the commonalities, and differences of language versus executive control impairments remain under-specified. We tailored a Bayesian hierarchical modeling solution in a largest-of-its-kind cohort (1080 stroke patients) to deconvolve language and executive control in the brain substrates of stroke insults. Four cognitive factors distinguished left- and right-hemispheric contributions to ischemic tissue lesion. One factor delineated language and general cognitive performance and was mainly associated with damage to left-hemispheric brain regions in the frontal and temporal cortex. A factor for executive control summarized control and visual-constructional abilities. This factor was strongly related to right-hemispheric brain damage of posterior regions in the occipital cortex. The interplay of language and executive control was reflected in two factors: executive speech functions and verbal memory. Impairments on both were mainly linked to left-hemispheric lesions. These findings shed light onto the causal implications of hemispheric specialization for cognition; and make steps towards subgroup-specific treatment protocols after stroke.

## Introduction

In our rapidly aging societies, stroke is now the leading cause of long-term disability worldwide, with 12.2 million new cases each year.^1^ Globally, one in four people will be affected by stroke in their lifetime. Stroke often severely affects cognition and can cause loss of language and executive functions.^2–4^ These cognitive faculties are crucial for interpersonal interaction in everyday life. Language is a key ability for communication uniquely developed in humans, including production and comprehension abilities.^5^ Aside from specific linguistic operations, efficient communication also requires controlled planning, focusing and flexible thinking. These mental skills are subsumed as executive functions and include inhibitory control, working memory and cognitive flexibility.^6^ Brain damage can severely affect executive functions, including the ability to complete basic and complex activities of daily living and participate in work, social, and leisure activities.^7^ Yet, despite the wide-ranging impact of stroke-induced dysfunction on the individual patient’s cognitive abilities and our society at large, it remains unclear how the neural networks for cognitive functions recover from tissue damage.^8^ A deeper look into the commonalities and differences of stroke-induced tissue dysfunctions leading to language versus executive control deficits would identify biologically valid subgroups, thereby paving the way for more accurate outcome predictions and better-targeted therapeutics in the future of precision medicine.^9,10^

Indeed, today’s cognitive therapy after stroke routinely ignores the individual topography of the original brain lesion.^11^ Such one-fits-all treatment is probably a culprit for the observation that there is substantial variability across patients in treatment success, leading to overall small effect sizes.^12,13^ The critical gap in our knowledge about the specificity of stroke-induced impairments on different cognitive functions may partly result from the fact that most previous studies focused on impairments in single cognitive domains derived from small patient cohorts.^14–16^ Such studies suggest that language impairments result from damage to key language regions in the left hemisphere, while executive deficits have been preferentially linked to lesions in the right hemisphere.^8,16^ This distinction ignores the functional interplay of both domains, especially under challenging conditions. Indeed, recent work provides clear clues that successful language recovery after stroke may extend beyond classically studied language regions: neuroimaging studies suggest that domain-specific recovery of language functions also includes recruitment of domain-general networks for cognitive control, such as the multiple-demand network.^17–20^ This implies that some patients with deficits in the language domain may particularly benefit from training of executive functions. For example, in case of prevailing verbal executive deficits, treatment may be most successful if centered on both executive and speech functions.

However, the precise contributions of domain-general regions to language processing remain largely obscure. Neuroscientists are still to clarify if these regions have compensatory potential after the loss of specific language functions or mainly subserve support processes such as cognitive control functions when processing difficulty is increased.^21,22^ If domain-general regions essentially contribute to language recovery, one may expect overlap in the lesion-deficit associations for language and executive control, especially for language operations with higher executive demands. This hypothesis is based on the observation that under challenging conditions (e.g., noisy environments or cognitive decline), language-related activity engages both language-related as well as executive control regions.^e.g.,23^

Yet, as a barrier to progress, previous studies neglected a comprehensive characterization across several key cognitive domains in a single clean analysis. Indeed, most existing stroke patient cohorts are under-phenotyped and do not typically lend insight into cognitive measures that subsume various domains derived from the same cohort. Consequently, commonalities and differences between cognitive domains are unclear. Moreover, only a few studies to date take a network perspective and consider the role of both hemispheres across various cognitive domains in patients with stroke.^e.g.,15,24,25^ While the few existing studies lend initial evidence for substantial lateralization differences between language and executive control, they do not address the functional interplay of domains.

Confronting these shortcomings, our present investigation was designed to tease apart the overlap and diverging consequences of lesion topologies for language and executive control in a richly phenotyped cohort of >1,000 patients with stroke. To draw a complete picture of major domains for human cognition, we examined multiple verbal and non-verbal assessments with varying executive demands. This comprehensive characterization, in combination with a multivariate factor analysis, allowed us to extract key dimensions of human cognition and identify the impact of stroke lesions on different regions of functional capacity.

Our first hypothesis was that lesions to the left-hemispheric language network (e.g., left prefrontal, anterior, and posterior temporal cortex^5,26^) would preferentially track language impairments. In contrast, disturbed executive control functions should index damage of the distributed (potentially right-dominant) multiple-demand network.^19,27,28^ Second, we expected a considerable degree of overlap of executive functions with the whole-brain lesion distribution for language assessments that require high control demands. These assessments should draw on both language-specific and domain-general control regions, with a potential strongest overlap in bilateral prefrontal control regions.^19,20,29,30^

To foreshadow the main results of our population-scale lesion-network investigation, we identified four factors which define unique left- and right-hemispheric contributions to different cognitive domains. One factor delineated language and general cognitive performance and was mainly associated with damaged regions in the left hemisphere. An executive control factor summarized control and visual-constructional abilities and was strongly related to right-hemispheric brain damage. The interplay of language and executive control was reflected in two different factors, delineating executive speech functions and verbal memory. Impairments on both were mainly linked to left-hemispheric lesions. Collectively, our findings formulate new insight into causal elements of structure-function underpinnings and hemispheric specialization in an approach that cuts across cognitive domains usually studied in isolation.

## Material and Methods

### Characteristics of the participant sample

Details on participant recruitment can be found in the **SI Materials and Methods** section. Most of the patients had lesions in either the left or right vascular territory of the middle cerebral artery and there was no significant difference in the number of lesioned voxels per patient between the left and right hemisphere (two-sided t-test: p=0.81). The maximum of lesioned tissue was localized in subcortical zones. Patients underwent a rich battery of neuropsychological tests ∼three□months after the acute onset of stroke (median time post-stroke: 98□days, SD: 66).^31^ Specifically, each patient was characterized by eight key assessments of post-stroke cognitive performance: global cognitive function (Korean version of the Mini Mental State Exam, MMSE)^32^, language (Korean short version of the Boston Naming Test, BNT)^33^, executive speech function (Korean version of phonemic and semantic fluency tests)^31^, executive control functions (Trail Making Test, TMT, version A and B and Digit Symbol Coding Task, DSCT)^34,35^, visuospatial functioning (Rey Complex Figure, RCF)^36^, as well as verbal learning and memory function (Soul Verbal Learning, SVL).^31^

Performance in the MMSE reflects global cognition, including the orientation to time and place, as well as calculation or language performance. The BNT (Korean short version) is a standardized clinical test that measures word retrieval of patients by asking them to name 15 pictured nouns (short version). Semantic and phonemic fluency tests are prototypical measures of verbal fluency, probing executive functioning, speed and attention, and access to the mental lexicon. In semantic fluency tests, subjects are required to generate words that belong to a specific category for a limited time window. Phonemic fluency tests require the subject to generate words starting with a given letter. The TMT probes visual attention and task switching in two parts in which the subject is instructed to connect a set of dots or numbers and letters as quickly as possible while maintaining accuracy. The test provides information about visual search speed, scanning and processing speed, mental flexibility, and executive functioning. The DSCT was designed to measure processing speed, working memory, visuospatial processing, and attention. In this test, subjects learn a code in which each digit is represented by a symbol. Subjects have to substitute the correct symbols for a series of digits as quickly and accurately as possible. In the RCF, patients are first asked to copy a complex line drawing and then draw it from memory. The test measures visuospatial constructional abilities and visual memory. The SVL, in turn, examines episodic memory performance and requires the auditory learning of a word list and tests its memorization by an immediate recall task.

Finally, the Informant Questionnaire on Cognitive Decline in the Elderly (IQCODE) was used to capture the cognitive performance before the ischemic event.^37^ IQCODE is an established retrospective assessment relying on health-proxy reports that provide a widely used and validated measurement instrument of cognitive decline in the ten years before stroke onset. All tests and questionnaires were conducted by a trained clinical psychometrician blinded to clinical or neuroradiological patient information.^38^ Additionally, we collected available sociodemographic and clinical information, such as age, sex, time since stroke onset (in days), years of education and lesion volume (grey and white matter). Each continuous (non-binary) variable was z-scored across all subjects to ensure comparability.

### Data processing

For details on the preprocessing, the neuroimaging protocols, as well as the extraction of target lesion load signals and deconvolution of lesion atoms (i.e., lesion patterns, see **Fig. 1**), please refer to the **SI Materials and Methods**.

**Figure 1.**
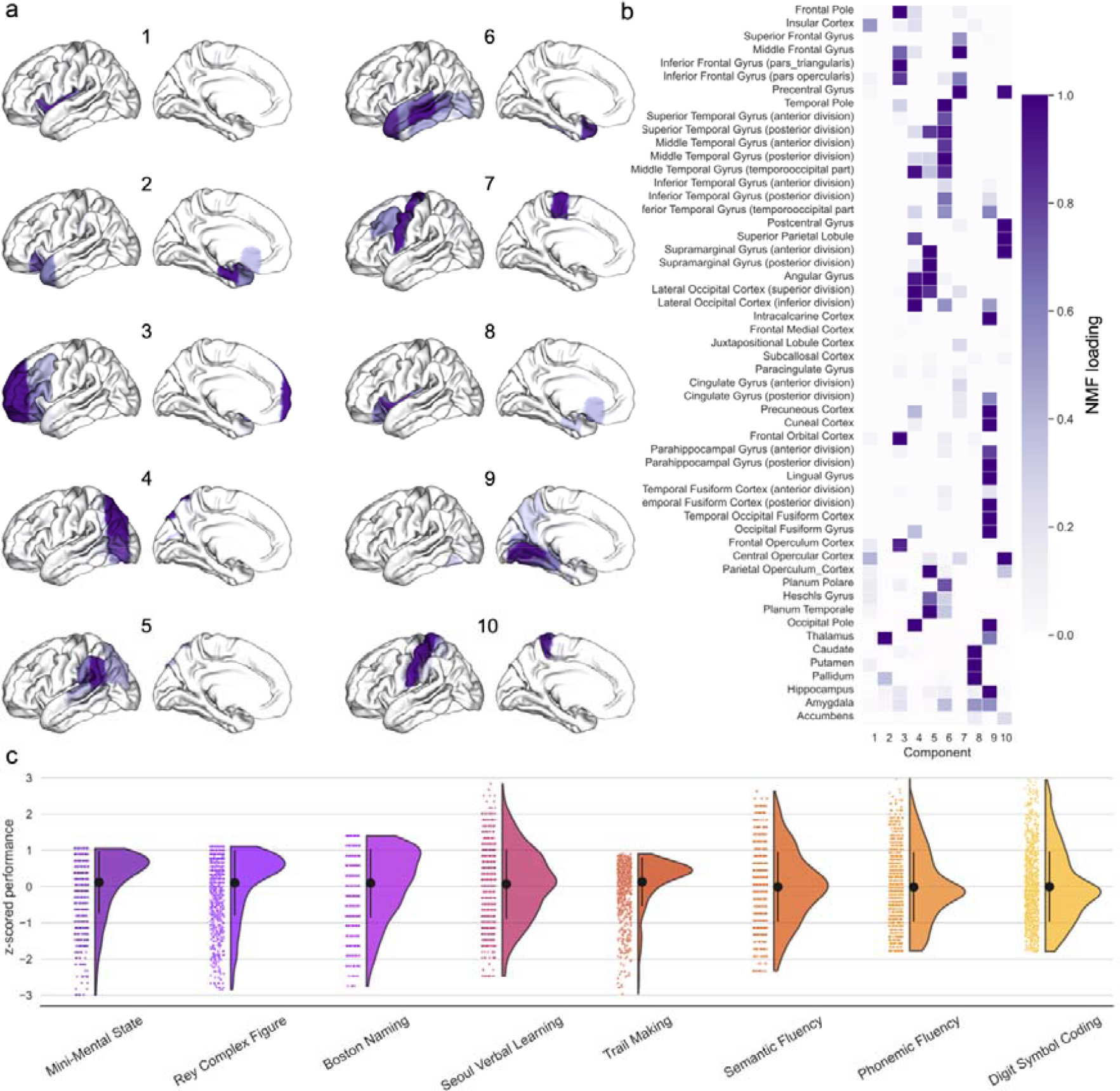
Lesion atoms of stroke patterns reveal unique lesion topologies across the whole brain. Voxel-wise information on stroke-induced lesions from >1,000 patients was summarized based on 108 anatomical region definitions from a reference atlas (54 per hemisphere) and derived by dimensionality-reducing pattern learning. Region-wise lesion measures were then compressed into 10 essential lesion-pattern “prototypes” in each hemisphere by capitalizing on non-negative matrix factorization (NMF). **(a)** Our derived 10 brain lesion atoms projected on the cortical surface. The resulting lesion atoms capture biologically plausible lesion pattern topographies. **(b)** Relevance of specific brain regions within each of the 10 lesion atoms (quantified as NMF loading) shows whole-brain coverage with distributed lesions in frontal, temporal and parietal as well as subcortical regions. Lesion atoms 1 and 2 implicate the insular cortex and thalamus, respectively. Lesion atom 3 covers the prefrontal cortex (including the inferior frontal gyrus). Occipital regions are covered by lesion atom 4 (inferior occipital cortex and adjacent inferior temporal and parietal regions) and 5 (superior occipital cortex and heteromodal association regions of the inferior parietal lobe, angular gyrus, supramarginal gyrus). Temporal regions are covered by lesion atoms 6 (superior, middle, and inferior temporal cortex) and 9 (fusiform gyrus, hippocampus and parahippocampal regions, as well as precuneus and cuneus in the parietal lobe). Precentral and postcentral regions are included in lesion atoms 7 and 10. Lesion atom 8 includes the basal ganglia. **(c)** Z-scored behavioral performance. Raincloud plots show the performance of each subject for the eight cognitive assessments. Our analyses benefited from a dedicated data-driven lesion mapping strategy tailored to deeply phenotyped patients with stroke.

### Latent factors driving cognitive outcomes

We explored the possibility that the observed inter-correlations between our target eight cognitive scores are influenced by one or more underlying factors that are not directly observable. This assumption is in direct analogy to the q-factor in intelligence research as well as the big-5 personality model, where factor analysis was applied to personality survey data to summarize aspects of personality into five broad dimensions.^39^ The factor-analysis-derived overarching domains contain and subsume the most known personality traits and are assumed to represent the basic structure behind all personality traits.^40^ Hence, we turned to factor analysis to uncover coherent latent patterns explaining the interrelationships among the set of our cognitive scores (see **SI Materials and Methods** for details).

### Predicting cognitive outcomes at 3 months post-stroke

The NMF-derived expressions of topographical lesion atoms provided the neurobiological input into our Bayesian hierarchical models.^41^ Carefully tailored to two different analytical strategies, we built two distinct classes of Bayesian hierarchical models. First, we designed a single multivariate, multivariable Bayesian hierarchical model to explain interindividual differences in eight cognitive outcome scores. The multivariable aspect allowed us to jointly estimate population means, variances and covariance of interrelated post-stroke outcomes. This model is labeled as MIMO (multiple-input multiple-output) throughout the manuscript. In the second analytical scenario, we derived a set of four multivariate Bayesian hierarchical models dedicated to the four factors representing investigated cognitive domains. In other words, each model distilled knowledge from multiple inputs to provide predictions for a single output (i.e., factor). This setup is referred to as MISO (multiple-input single-output) throughout the manuscript. Finally, all models took into account several covariates, including age, age^2^, sex, time since stroke onset, education years, pre-morbid cognitive performance and total lesion volume following previous research.^24,25^ Details on the Bayesian model specification and posterior Bayesian estimates for all parameters and models are available in the **SI Materials and Methods** section.

Our Bayesian hierarchical approach facilitated the careful dissection of predictive relevance allocated to different levels of the model. For all outcome models, we first evaluated lateralization effects inferred from the left and right hemisphere posterior dispersion distributions. Subsequently, we considered the lateralization effects of specific lesion atoms and, lastly, reverted back the predictive relevance of lesion atoms^42^ to the level of the anatomical brain regions for each modeled outcome. Finally, we inspected covariates in conjunction with hemisphere or lesion atom contributions to concurrently quantify the contribution of a lesion representation to the outcome while rendering explicit the influence of important demographic, sociodemographic and clinical characteristics.

### Quantifying the similarity with language and multiple-demand network

We explored the similarity of our factor-specific brain maps of beta coefficients with maps of the language network and domain-general multiple-demand network.^43,44^ Both maps are freely available online (https://evlab.mit.edu/funcloc/). As a first step, for each factor-specific Bayesian model, we inverse-projected beta coefficients from the latent space of non-negative matrix factorization to the space of the Harvard-Oxford atlas. Thus, we obtained a coefficient for each region of this atlas. Then, we projected the absolute values of region-specific coefficients on the cortex. Finally, to quantify the similarity, we computed Pearson’s r using *neuromaps* toolbox^45^ between the brain map of coefficients pertinent to a given factor and brain maps of language as well as multiple-demand networks.

### Data and code availability

Anonymized data that support the findings of this study are available from the corresponding author DB upon reasonable request. Analyses were conducted in a Python 3.7 environment and predominantly relied on the packages nilearn and pymc3. Full code is accessible to and open for reuse by the reader here: https://github.com/jakubkopal/bayesian_stroke.

## Results

Our study set out to make steps towards clinically useful single-patient predictions of language-specific vs. domain-general executive functions. We first derived a whole-brain representation of distributed lesion patterns from expert-segmented post-stroke lesion maps. Next, we devised and deployed a principled Bayesian framework. A fully probabilistic framework is well suited to trace out how brain tissue impairments impact interindividual differences in key cognitive domains after acute ischemic stroke^24,25^ and carefully quantify the extent of overlap of these brain-behavior correspondences. The combination of a comprehensive cognitive test battery with factor analysis in a Bayesian modeling framework should capture the impact of stroke-induced dysfunction in distributed brain regions for key faculties of human cognition.

### Cognitive target endpoints of a deeply phenotyped patient cohort

We analyzed information from structural brain scans of 1080 ischaemic patients with stroke obtained from a prospective South-Korean stroke registry.^24,31^ MRI acquisition was performed within one week after the stroke to predict cognitive outcomes three months after stroke. Details about structural brain image assessment and resulting segmented lesion patterns were described elsewhere.^24,25^ Neuropsychological and demographic data were obtained from the Korean-Vascular Cognitive Impairment Harmonization Standards-Neuropsychology Protocol.^31,46^ Our analyses focused on established assessments of global cognitive function (MMSE)^32^, language (Korean short version of the BNT)^33^, executive speech function (Korean version of phonemic and semantic fluency tests)^31^, executive control functions (TMT, version A and B and DSCT)^34,35^, visuospatial functioning (RCF)^36^, as well as verbal learning and memory function (SVL)^32^, thus indexing a spectrum of language production and executive control functions in each patient.

### Stroke lesion atoms isolate a typology of unique lesion constellations

We first summarized the high-dimensional lesion information at the voxel level in 108 cortical and subcortical brain regions (54 per hemisphere, based on the Harvard-Oxford cortical atlas). We then aimed to uncover coherent hidden patterns of lesion topography directly from the brain scan data themselves. To this end, we applied a previously established set of 10 distinct topographical lesion configurations, so-called “lesion atoms” (lesion patterns), in a data-driven fashion based on non-negative matrix factorization (NMF).^24,25,47^ Note that the definition of lesion patterns was symmetrically identical in both hemispheres. The spatial definition of the atomic lesion patterns corresponded to biologically plausible components of stroke-induced brain lesions in both hemispheres and their co-occurrences across patients (**Fig. 1a**). Accordingly, lesion atoms were reminiscent of territories of arterial blood supply via the anterior (lesion atoms 3 and 7), middle (cortical: 6, 7, 10, subcortical: 1, 8) and posterior (2, 4, 5, 9) cerebral artery territories. Important to our present analyses, our collection of lesion atoms showed whole-brain coverage with distributed lesions in frontal, temporal and parietal as well as subcortical regions (**Fig. 1b** for details).

Having parsed the lesion-segmented brain scans by means of the recurring stroke topographies, we investigated the overlap versus dissociation of lesion patterns on cognitive outcomes after stroke (**Fig. 1c** for the distribution of cognitive scores).

### Bayesian modeling predicts atom-impairment-links in a hemisphere-aware fashion

Next, we performed a Bayesian analysis to test how the patients’ configurations of lesion atoms explain interindividual differences in post-stroke outcomes. Since these post-stroke outcomes are expected to be interrelated, we derived a single multiple-input multiple-output (MIMO) model. The MIMO model specification allowed us to jointly estimate population means, variances and covariance of our collection of eight outcome measures. This analysis can infer accurate uncertainties for the contribution of each hemisphere and their dependent lesion atom to the successful endpoint prediction at the single-patient level. Our brain measurements were modeled by taking into account potential sources of variation outside of primary scientific interest: total lesion volume, age, age^2^, sex, education in years, and premorbid cognitive performance^24,25^ (cf. methods). As a litmus test of model quality, the posterior predictive checks confirmed the adequate fit of our estimated MIMO prediction model. The variation was described by how individual lesion atom expressions explained differences in cognitive performance outcomes in patients^48^, albeit with different achieved performances across the target cognitive scores (coefficient of determination R^2^ ranged from 0.11 to 0.49 for phonemic fluency and MMSE, respectively, **Fig. 2A**). This empirical verification of the MIMO model’s ability for brain-based predictions of post-stroke capacities is a well-recognized proxy of external validation given the actual data at hand.^49^ The estimated standard deviation posteriors captured the overall left and right hemispheric relevance with good certainty (i.e., the width of the posterior variance parameter distributions) and dissociated hemisphere-specific predictive contributions tiled across all candidate lesion atoms (**Fig. 2B**).

**Figure 2.**
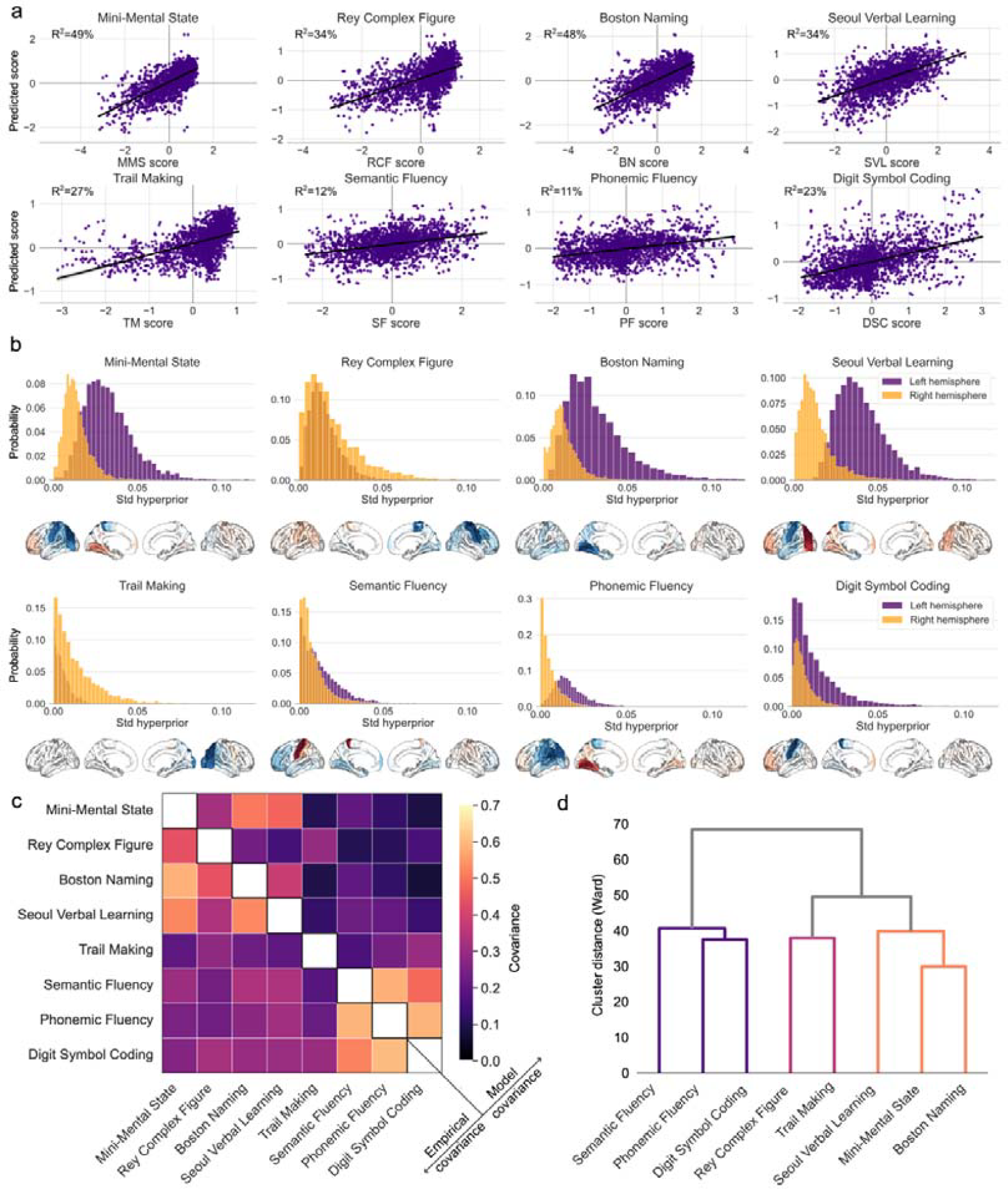
Bayesian hemisphere-aware analysis can robustly predict individual clinical outcomes. **(a)** Model performance of the inferred MIMO (multiple-input multiple-output) Bayesian analytical solution using brain lesion atoms in predicting cognitive impairments. Posterior predictive checks were performed for the Bayesian models that were estimated to predict interindividual differences in eight cognitive measures (on z-scale). Model-based simulations (y-axis) were compared to the observed data (x-axis) to compute the overall explained variance (coefficient of determination, R^2^). **(b)** Posterior probabilities for the hemisphere-specific lateralization effects for each clinical endpoint. Lesions to the left hemisphere were more relevant for single-patient predictions of cognitive outcomes in the MMSE, BNT, SVL semantic and phonemic fluency tests, and DSCT. In contrast, damage in the right hemisphere was more explanatory for performance in the RCF and TMT. The posterior model parameters correspond to the upper hemisphere level of our Bayesian multilevel modeling strategy that uncovered the hemisphere-specific model certainty for each cognitive performance dimension. Surface projections: blue colors: lesions are associated with relatively stronger impairments, red colors: lesions are associated with relatively weaker impairments. **(c)** The covariance matrix quantifies the co-occurrences between cognitive outcome measures across patients. Covariance is high (>0.5) for MMSE, BNT and SVL, as well as for both fluency measures and DSCT. In comparison to empirical covariance, the model covariance matrix quantifies the intrinsic relationship between cognitive outcomes given brain lesion conditions and other covariates. **(d)** Similarity of brain-behavior representations: Hierarchical clustering of model-specific coefficients obtained per clinical endpoint shows distances (similarities) among cognitive measures based on Ward’s method. A first cluster summarizes both fluency assessments and DSCT. Based on distances, RCF and TMT can also be grouped. Finally, SVL, MMSE and BNT show low distances, suggesting high similarity. Overall, our hemisphere-aware Bayesian model revealed differences in cognitive outcome predictions that follow the distinction between measures of language and executive function.

As a central feature of our findings, lesions in both hemispheres contributed in specific amounts to prediction success across the examined cognitive endpoints. Lesions in the left hemisphere were overall more relevant for single-patient predictions of most cognitive outcomes. More specifically, the estimated model parameters for hemisphere-specific dispersion showed relative left-biased lateralization effects for MMSE, BNT, fluency tests, SVL and DSCT; but relative right-biased lateralization for RCF and TMT. These findings were in line with the empirical and model-derived covariance matrices between cognitive outcome measures which also revealed strong similarity between MMSE, BNT and SVL, as well as between both fluency measures and DSCT (**Fig. 2c**). Finally, we used hierarchical clustering of the derived beta estimates (corresponding to lesion atom involvements from the estimated MISO prediction model) to explore similarities between cognitive outcome tests (**Fig. 2d**). This post-hoc analysis showed similarities for both fluency assessments and DSCT. RCF and TMT were also close to each other in this clustering space. The last cluster based on the Ward similarity measure was formed by SVL, MMSE and BNT. Collectively, our Bayesian approach revealed hemisphere-specific differences between cognitive measures based on predictions from lesion topographies. Our observations suggested that some of the cognitive measures can be considered as representative cognitive key dimensions.

### Four impairment dimensions reveal hemisphere-specific effects on language and executive functions

We performed an exploratory factor analysis to extract principled representations of cognitive domains with the possibility to cut across our eight outcome measures. Based on the scree test criterion, we favored a four-factor solution of overarching cognitive dimensions (**Fig. 3a**). The four-factor solution provided a faithful characterization of the overarching cognitive dimensions that most effectively summarized the behavioral variation across eight clinical endpoint assessments.

**Figure 3.**
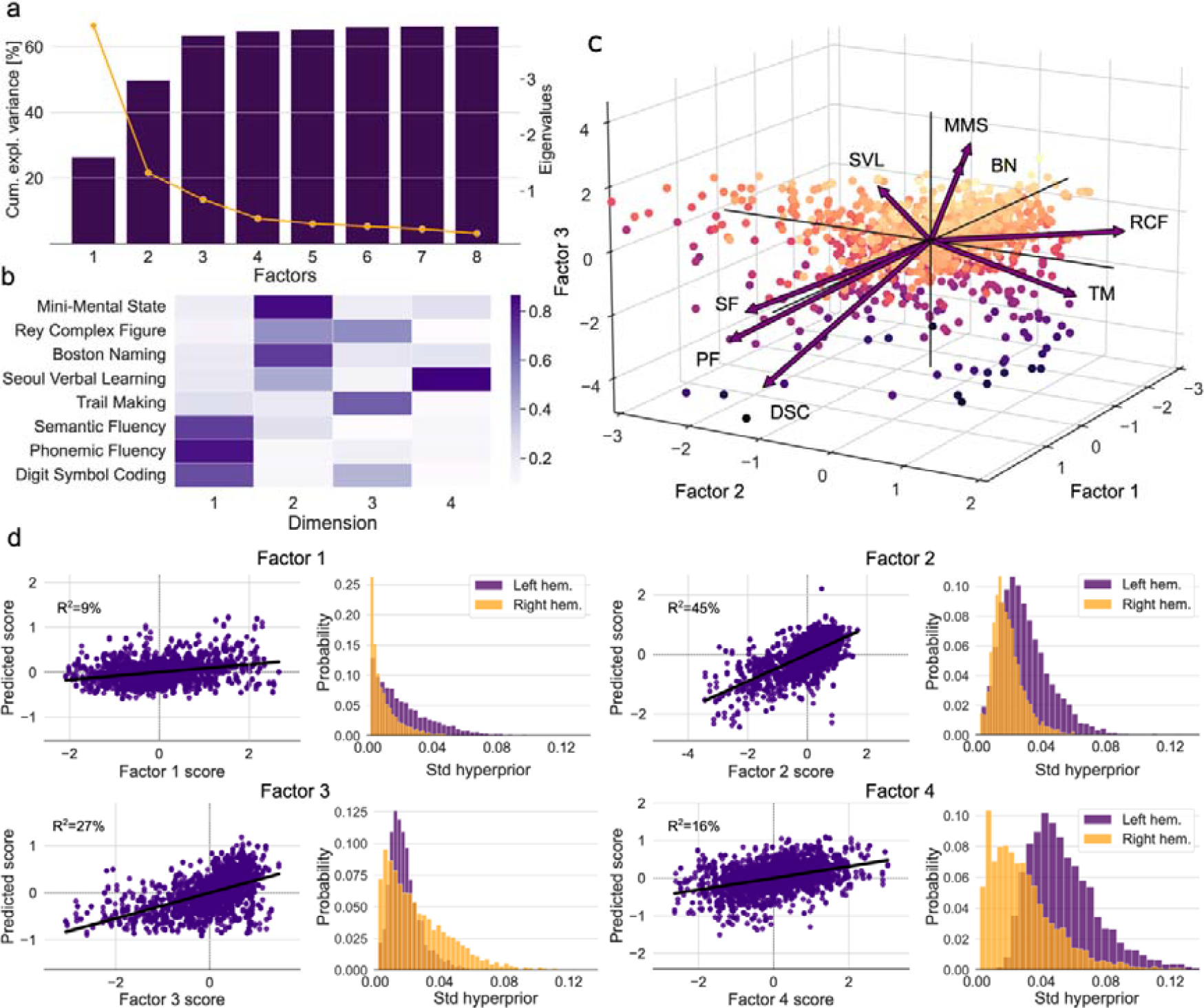
Four factors trace out the similarity and dissimilarity between language-related and executive function-related clinical endpoints. Low-dimensional representation of clinical endpoints dissociates between language-related and executive control-related factors. **(a)** Scree plot illustrating the amount of explained variance in outcome measures and associated eigenvalues. According to the scree test criterion, a four-factor solution is an effective low-dimensional data representation which explains a substantial portion of the variation. **(b)** Factor loadings for the included outcome measures. **(c)** Factor biplot depicting participant scores and loading vectors for the first three factors. Loading vectors represent how strongly each characteristic influences the resulting factor. The angle between a pair of vectors corresponds to the correlation between the given characteristics. Participant factor scores are displayed as points in the plane formed by three principal components. Dot colors code for factor scores (bright color: high score). **(d)** Performance of the inferred MISO (multiple-input single-output) Bayesian analytical solutions in predicting cognitive impairments for each factor. Posterior predictive checks are shown for the Bayesian models that were estimated to predict interindividual differences in (z-scored) factors. Model-based simulations (y-axis) were compared to the observed data (x-axis) to compute the overall explained variance (coefficient of determination, R^2^). Right panels: Relevance of the posterior parameter distributions (std=standard deviation) of the left and right hemisphere obtained through separate Bayesian hierarchical models dedicated to predicting each factor. Lesions to the left hemisphere were more relevant for single-patient predictions of cognitive outcomes for factors 1, 2 and 4. In contrast, lesions in the right hemisphere were more relevant for outcome predictions for factor 3. The posterior model parameters correspond to the upper hemisphere level of our Bayesian multilevel modeling strategy that uncovered the hemisphere-specific model certainty for each factor. The four-factor solution, in combination with four Bayesian models, successfully untangled the unique contributions of cognitive outcomes along the language vs executive control axis.

Factor 1 was labeled as **executive speech functions** because it explained the combined variation of phonemic and semantic fluency and DSCT in our patients. Factor 2 tracked the overlap in the **language and general cognitive performance** by explaining a notable variation of MMSE and BNT, with smaller contributions estimated for RCF and SVL. Factor 3 captured **executive control functions**, as shown by the highest loadings of TMT and lower contributions of RCF. Finally, factor 4 represented **verbal memory** with high loadings of SVL. The four-factor solution (**Fig. 3b,c**) supported and complemented the results from the hierarchical clustering approach (**Fig. 2d**). Specifically, the four-factor solution revealed that some impairment dimensions of language and executive functions are clearly distinguishable, while there is a shared core in the dimension of executive speech functions and verbal memory.

To explore the brain lesion implications of the derived four intrinsic cognitive factors, we estimated a set of four Bayesian hierarchical models. This multiple-input single-output (MISO) model class was carried out so that each model was dedicated to a single factor. The probabilistic prediction of patient-specific factor expressions across the four MISO models achieved 9-45% explained variance (R^2^, coefficient of determination; factor 1: 9%, factor 2: 45%, factor 3: 27%, factor 4: 16%). In line with the modeling results for the single outcome measures, factors 1, 2 and 4 were left-lateralized, whereas factor 3 showed right-hemispheric lateralization (**Fig. 3d**). Collectively, the four-factor solution in combination with four dedicated Bayesian MISO models successfully untangled the unique regional brain lesion contributions of cognitive outcomes along the language vs executive control axis.

### Cognitive factors associate with distinct lesion patterns in specific brain regions

Next, we explored how the derived four MISO models tied into four cognitive factors (i.e., cognitive symptom sets that vary across patients) are associated with specific atlas regions (**Fig. 4**) and particular lesion atoms (**Fig. 5**, see next section). For cognitive factor 1 (indexing primarily verbal fluency and executive control), we found that lesions to the left putamen, precentral gyrus, caudate nucleus, posterior superior and middle temporal gyrus (STG and MTG), and supramarginal gyrus were driving deficits in executive speech functions. In contrast, lesion-deficit associations for cognitive factor 2 (particularly reflecting word retrieval and overall cognitive abilities) were located more posterior, with the strongest impairments in language and general cognitive performance caused by lesions to the left lateral occipital regions, lingual gyrus, hippocampus and occipital-fusiform gyrus. Complementing the pattern for factor 2, cognitive impairments in factor 3 (indexing primarily executive control) showed prominent associations of executive deficits with right occipital regions, as well as the postcentral gyrus, superior parietal lobe and occipital-fusiform gyrus. For factor 4 (indexing primarily verbal learning and memory), we found relevant associations between cognitive deficits in verbal memory and lesions located in the left post- and precentral somatomotor regions, while damage to the left and right occipital regions was associated with preserved functions. Overall, the derived cognitive impairment dimensions revealed distinct ties to brain lesion patterns in both hemispheres, following a left-versus-right and anterior-posterior axis: impairments in executive speech functions were associated with left-hemispheric damage to subcortical, precentral, and more anterior temporal regions. Deficits in general cognitive and language abilities were related to more posterior damage, including in the left occipital and temporal regions. Impaired executive functions were associated with lesions in the right occipital and temporal regions. Finally, deficits in verbal memory were related to lesions in the left postcentral regions. The overall distribution of lesion patterns across factors showed that language and executive functions mainly dissociate in posterior regions (including occipital, posterior temporal, and subcortical regions), with language being located relatively more anterior. In contrast, the overlap of both functions was found in more anterior regions, including pre- and postcentral somatomotor gyri of the cerebral cortex.

**Figure 4.**
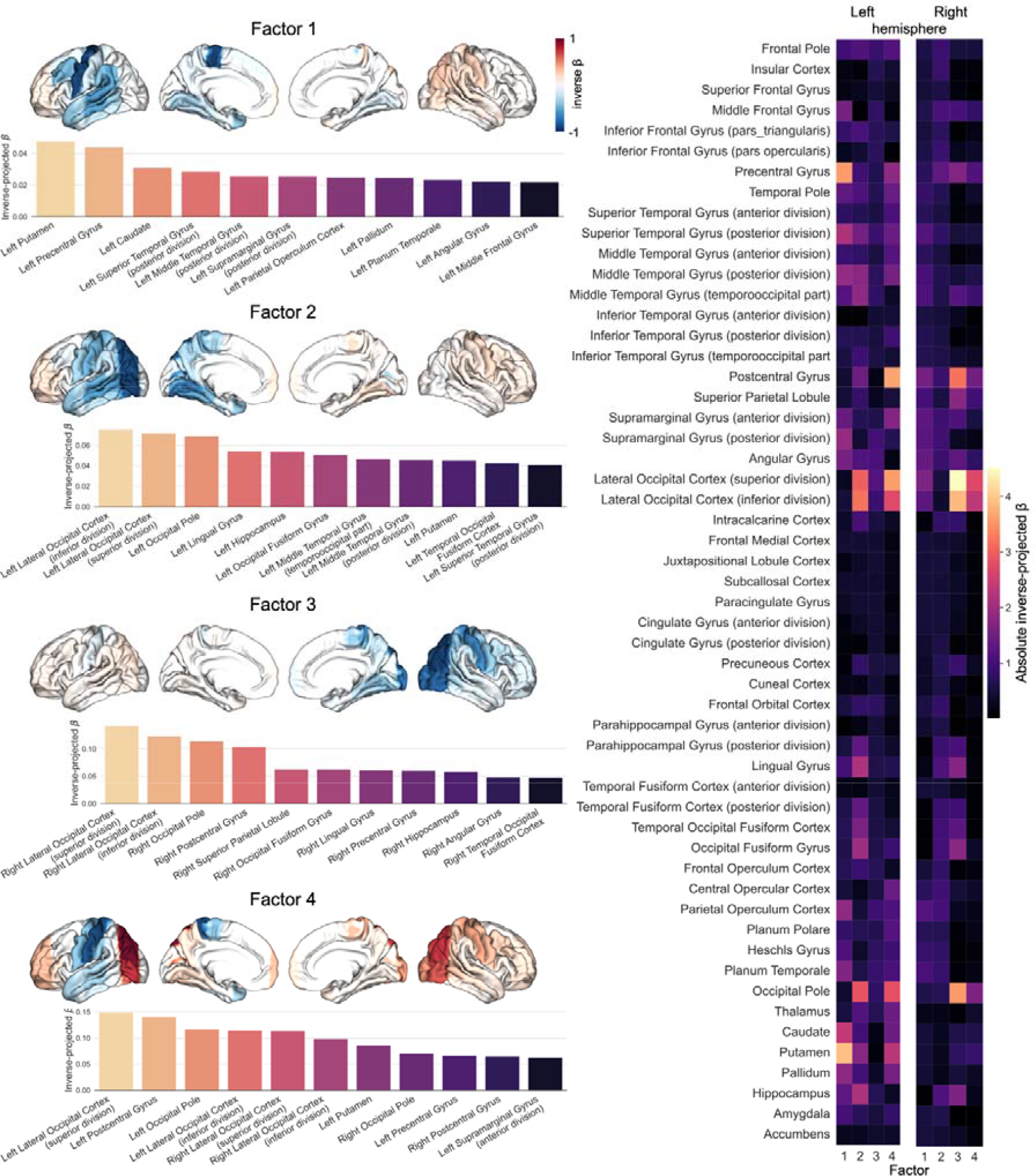
Four factor-specific models trace out distinct stroke-induced lesion patterns for single regions across different cognitive dimensions. Left side: Lesion-deficit prediction for each of the four factors. Colored brains reflect associations of brain regions with lost (blue color) or preserved (red color) cognitive functions. Note that factors 1, 2, and 4 show stronger left-hemispheric lateralisation, while factor 3 is right lateralized. Right side: 54 parcels per hemisphere were included based on the Harvard-Oxford cortical and subcortical atlas. The predictions of four factors with differential contributions to language vs executive functions are subserved by distinct brain patterns.

**Figure 5.**
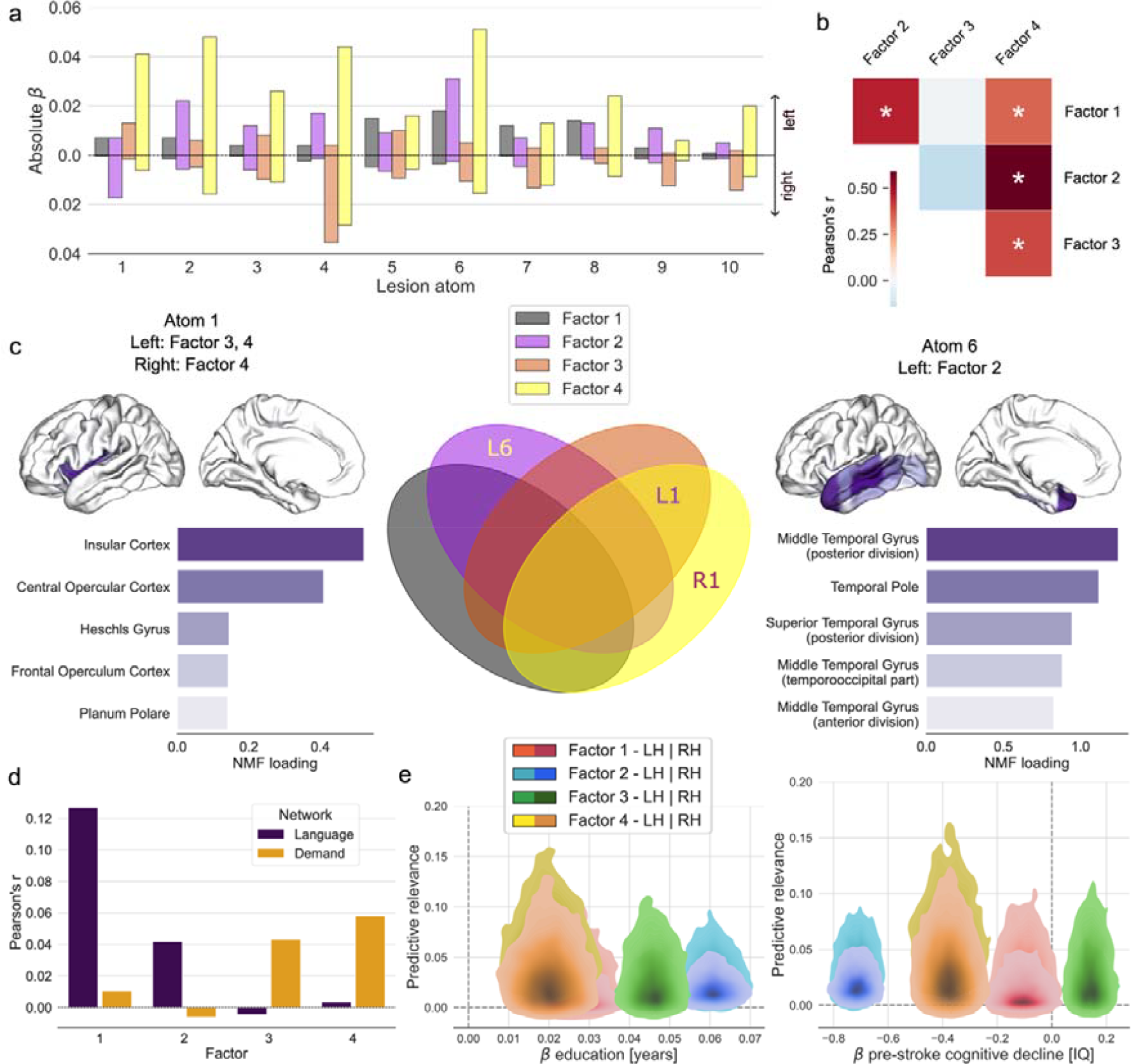
Distinct associations of lesion atoms and cognitive factors disentangle language and executive deficits. Results for the four-factor solution. **(a)** Lesion-deficit associations for the different factors and lesion atoms. Factors 1, 2 and 4 show the most prominent brain lesion effects for atom 6 in the left hemisphere (including temporal regions), while factor 3 shows strong implications for atom 4 (summarizing occipital regions) in the right hemisphere. **(b)** Correlations between model-specific beta values dedicated to each factor. Left-dominant factors show more significant correlations with each other. **(c)** Distinct relationships between significant lesion atoms and cognitive impairments. The significance of lesion-related beta estimates was assessed based on the difference of 90% highest density intervals from zero. Left: lesion atom 1 is characterized by left-hemispheric contributions from factors 3 and 4 and right-hemispheric contributions from factor 4, with the overall strongest load of the left insular cortex. Middle: Overlap and distinct contributions of the four factors. Right: lesion atom 6 is characterized by left-hemispheric contributions from factor 2, with the strongest load of the middle temporal gyrus. **(d)** Correlations of factors with maps of the language network and multiple-demand network^44,45^ show a relatively stronger overlap of executive speech functions and language with the language network. Executive functions and verbal memory show relatively stronger associations with the multiple-demand network. **(e)** Hemispheric relevance depends on key covariates. The plot depicts the interrelation between the relevance of lesion load in each hemisphere (y-axis, left hemisphere: light colors, right hemisphere: dark colors) and marginal posterior parameters of two key covariates (x-axis) in predicting cognitive performance. Years of education had positive effects on all four factors. The strongest effect was on factor 2. Conversely, an increase on the IQCODE scale^37^, i.e. a higher pre-stroke cognitive decline, predicted a decrease in scores of factors 1, 2, and 4. Lesion atom-outcome associations uncovered factor-specific language and executive control deficits: a prominent contribution of the lesion atom covering left-hemispheric superior and middle temporal regions to language, executive speech functions and verbal memory and strong implications of right occipital brain regions for executive functions.

### Lesion atom-outcome associations uncover factor-specific language and executive control deficits

Finally, we complemented region-based lesion-deficit associations with pattern-level results based on the identified lesion atoms. Factors 1, 2 and 4 showed the most prominent brain lesion effects for atom 6 (including inferior, middle, and superior temporal regions) in the left hemisphere (**Fig. 5a**). In contrast, factor 3 featured the strongest brain effects for atom 4 (including occipital and adjacent inferior temporal and parietal regions) in the right hemisphere. Additionally, factor 1 showed a robust implication of tissue lesions falling into the territories of lesion atoms 5, 7 and 8 in the left hemisphere (covering the superior occipital cortex and adjacent inferior parietal regions, precentral cortex and adjacent middle as well as inferior frontal regions, and basal ganglia). Factor 2 showed additional effects for atoms 2, 3, 4 and 8 in the left hemisphere (covering thalamus, (pre-)frontal cortex, occipital cortex and basal ganglia) and atom 1 in the right hemisphere (insular cortex). Factor 3 had additional implications of atom 1 in the left hemisphere (insular cortex) and atoms 6, 7, 9 and 10 in the right hemisphere (covering temporal regions, precentral, inferior and middle frontal regions, (para-)hippocampus and adjacent fusiform gyrus, pre- and postcentral cortex as well as the superior parietal cortex). Finally, factor 4 was additionally associated with brain lesion effects captured by all atoms except atom 9 in the left hemisphere and atoms 2, 4, 6 and 7 in the right hemisphere (thalamus, occipital cortex, distributed temporal regions, precentral and middle frontal gyrus). Overall, these results showed that stroke-induced language deficits could be mainly characterized by left-hemispheric lesions to prefrontal and temporal regions, while executive deficits are related to right-hemispheric damage in occipital and postcentral regions. Impairments in the overlap of both domains, summarized as executive speech functions and verbal memory, were associated with more distributed lesions in the left precentral and prefrontal regions as well as posterior temporal and parietal regions.

Notably, we found correlations between factors with left-hemispheric dominant lesion-outcome associations (factors 1, 2, 4, **Fig. 5b**). Factors 3 and 4 were also highly correlated. Next, we investigated differences in the significant contribution of specific lesion atoms. In other words, we probed the significance of each beta estimate by assessing whether the underlying 90% highest density interval included zero. Significant associations between deficits and lesion atoms were found for atoms 1 and 6 (**Fig. 5c**). Lesion atom 1 (highlighting insular cortex) showed left-hemispheric contributions from factors 3 and 4 and right-hemispheric contributions from factor 4. Lesion atom 6 (temporal lobe regions) was characterized by left-hemispheric contributions from factor 2.

Correlating our factors with maps of the language network and domain-general multiple-demand network^43,44^ revealed a relatively stronger overlap with the language network for factors 1 and 2 (executive speech functions and language). In contrast, factors 3 (executive functions) and 4 (verbal memory) showed more overlap with the multiple-demand network (**Fig. 5d**). We also explored interrelations between important covariates (i.e., age and years of education, as well as pre-stroke cognitive decline and number of lesions) in the prediction of cognitive outcomes for each factor **(Fig. 5e)**. While we observed a positive impact of education on cognition-related factor scores, the inverse was true for cognitive decline.

Collectively, our results showed unique hemisphere-specific contributions of language and executive functions. These specific contributions dissociated at the level of brain regions (including prominent contributions of temporal, (pre-)frontal and occipital regions) and networks. Aggregated cognitive factors derived from the factor analysis showed stronger associations of executive speech functions (factor 1) as well as language and general cognitive abilities (factor 2) with the previously described language network^42^. In contrast, executive control functions (factor 3) and verbal memory (factor 4) were relatively more strongly associated with the multiple-demand network.^49^ While factors 1, 2 and 4 showed prominent associations with left-hemispheric lesion atoms, factor 4 showed additional brain effects in right-hemispheric lesion atoms, and factor 3 revealed a prominent association with right-hemispheric lesion atoms. The observed constellations of specific and overlapping associations helped to uncover unique and general contributions and argue for a complex interplay of specialized and domain-general networks and brain regions across different key domains of human cognition.

## Discussion

Here, we shed light on the causal relevance of distributed brain regions realizing the cognitive axis of language versus executive control functions. We combined Bayesian hierarchical modeling with a rich portfolio of neuropsychological outcome measures in a large cohort of >1,000 patients with stroke. Seizing this opportunity, we were able to robustly predict individual clinical outcomes based on specific tissue lesion topologies. We identified four distinct cognitive factors that characterized the similarity and dissimilarity between language-related and executive control-related clinical outcomes. Our results elucidate specific lesion topologies associated with deficits in each of the two cognitive domains and detail their functional interplay.

As a core finding, our analytical framework could disentangle anterior-posterior and left-right principles for impairment of language and executive functions due to distributed stroke insults. Language and general cognitive performance impairments were primarily linked to lesions of prefrontal and temporal regions in the left hemisphere, while executive control was preferentially affected by damage of occipital and postcentral regions in the right hemisphere. As a second main finding, the interplay of both cognitive dimensions was reflected in two distinct factors, tracking executive speech functions and verbal memory capacities. Contrasting our expectations, impairments on these two cognitive factors were mainly linked to left-hemispheric lesion atoms, with only minor contributions of right-hemispheric regions. These identified left-hemispheric regions covered pre- and postcentral regions associated with cognitive control as well as motor cortices. Zooming in on specific lesion topologies across the whole brain provided a fine-grained picture of impairment dimensions associated with distinct lesion atom patterns and specific brain regions for each identified cognitive factor.

Overall, our findings illustrate that stroke-induced lesions in the left hemisphere were relatively more relevant for single-patient predictions of most examined cognitive outcomes. Accordingly, factors 1, 2 and 4 showed left-hemispheric dominance of lesion-outcome associations with the most prominent brain lesion effects for atom 6 in the left hemisphere. This lesion atom tracked grey matter lesions across the superior, middle, and inferior temporal brain regions, which are associated with language operations and global cognitive functions in studies of healthy and lesioned brains.^5,50–53^ The observed dominance of the left hemisphere in our findings may reflect our selection of cognitive outcome measures with an excellent coverage of verbal memory as well as speech and language-related faculties (i.e., MMSE, BNT, verbal fluency, SVL and DSCT).

Nevertheless, regarding the lesion patterns for the executive control-related factor 3, we were also able to isolate specific contributions of right-hemispheric regions. Moving beyond incumbent narratives from previous stroke research, our results further show that the shared substrates of language and executive control functions, that is, executive speech functions and verbal memory, mainly suffer from distributed left-hemispheric damage. Associated lesion atoms included both language regions as well as pre- and postcentral motor and control regions. In this context, we wish to highlight that relative to classical voxel-wise lesion symptom mapping approaches, our patient-specific predictions are better suited to forecasting clinical endpoints.^24^

Accordingly, hierarchical clustering and exploratory factor analyses quantitatively dissected the underlying cognitive dimensions in both hemispheres. The resulting factors showed links to specific lesion-deficit associations, establishing an anterior-to-posterior axis for executive speech versus language operations. The first factor summarized executive speech functions. Impairment of this cognitive factor was linked to lesions in distributed left-hemispheric brain regions. These included posterior and middle temporal as well as inferior parietal regions, which are frequently associated with phonological and verbal working memory processes (superior temporal gyrus and supramarginal gyrus^54–57^) or semantic control processes (middle temporal gyrus^58,59^). These neural processes are known to be relevant for verbal fluency. Other regions comprised left precentral and subcortical regions in the basal ganglia (putamen and caudate nucleus), linked to sequencing operations during speech processing and often reported in fluency tasks in the healthy brain.^60,61^ While the contribution of subcortical regions beyond the thalamus to clinical language impairments is still debated^62,63^, our in-depth quantifications add evidence for a key contribution of the basal ganglia in the left hemisphere to fluent speech operations. Notably, our findings in single brain regions were supported by complementary information from lesion atom-deficit associations, providing evidence for the relevance of the left precentral cortex and adjacent middle and inferior frontal regions, which are key regions for executive semantic processes^30,58,64^ and implicated in executive speech control.^66^ These results open a new window into how executive control is necessary for fluent language performance. Strikingly, the overlap in the lesion patterns shows that rather than recruiting additional control regions in the opposite hemisphere, fluent speech processing mainly draws on distinct left hemispheric regions, with a strong emphasis on the prefrontal and temporal cortex. This insight paves the way for future individualized lesion- and deficit-oriented therapeutic approaches that should include training of domain-general executive control elements for patients with impaired speech fluency.

The second factor particularly prioritized interindividual differences in language and general cognitive functions. Impairments were linked to lesions located in more posterior temporal regions of the left hemisphere which are associated with general cognitive abilities, as well as language and semantic memory.^66–69^ These regions included lateral occipital regions, lingual gyrus, hippocampus, and occipital-fusiform gyrus. Results from lesion atoms supported these findings and emphasized the role of left temporal and frontal regions for language and general cognitive abilities.^(see also 24)^ Collectively, the results for factor 2 highlight a strong contribution of lateral temporo-occipital regions to general cognitive and language functions associated with word retrieval.

The third factor carefully separated the stroke symptom component indexing central executive control, summarizing two measures (TMT and RCF) that are commonly considered to reflect executive control and visual-constructional abilities.^35,70–73^ A shared underlying cognitive dimension for both measures is supported by the observed similarities in our hierarchical clustering analysis. Cognitive impairments in factor 3 showed prominent associations of deficits in executive control functions with right occipital regions, as well as the postcentral gyrus, superior parietal lobe and occipital-fusiform gyrus. Accordingly, factor 3 showed strong implications for lesion atom 4, encompassing occipital and adjacent parietal regions in the right hemisphere. This lesion-deficit pattern likely reflects the strong reliance of the underlying tasks (TMT and RCF) on visual scanning and visual-constructional abilities.^72,74^ We note that the observed differences in the lesion-deficits patterns between factors 1 and 3 offer a fine-grained parcellation of the neural proxies of executive control functions. This implies that visuo-constructional abilities draw more strongly on posterior right hemispheric regions while verbal executive control is primarily related to left prefrontal and temporal regions. Collectively, these results move beyond a classic “left hemisphere = language, right hemisphere = executive functions’’ distinction, arguing for a key contribution of left hemispheric regions to verbal executive functions.

The fourth factor described verbal learning and memory performance in our patients, emphasizing the SVL task. This specific facet of cognitive impairment constellations was tied to damage in the left post- and precentral motor regions. Complementary evidence from lesion atom-deficit associations showed contributions of distributed regions in both hemispheres, likely reflecting the domain-general aspect of the SVL, drawing on verbal learning and memory functions.^31,76^ We note a close relationship of factor 4 with the neuropsychological assessments characterizing factor 2, as reflected in relatively high loadings of the SVL on factor 2, the strong positive correlation between factor 2 and 4, and the similarities between the SVL and the MMSE and BNT in the hierarchical clustering analysis. These results show that verbal memory is a unique cognitive faculty that quantifies the memory-related intersection of language and executive control functions, which can be clearly distinguished from executive speech functions as reflected in factor 1.

Finally, correlations with well-described brain networks for language and domain-general cognitive control^43,44^ provided complementary insight into lesion-deficit topologies. The two speech and language-related factors 1 and 2 showed positive correlations with the language network and negative correlations with the domain-general MDN. While this result was expected for the language-related factor 2, the specific overlap for factor 1 with language rather than MDN regions is surprising. This finding implies that executive speech functions may draw more strongly on language-related control processes rather than domain-general control functions associated with the MDN. This interpretation is supported by previous work arguing for a strong contribution of left prefrontal and posterior middle temporal regions to semantic control processes^30,58,76^, which are also necessary for accurate semantic fluency performance.^65,77^ The remaining factors 3 and 4 showed strong positive correlations with the MDN. The strong association of the right-dominant executive control factor 3 with this domain-general network was expected.^8,16,24^ In contrast, a strong association of the left-dominant factor 4 with the MDN may be surprising at first glance but likely reflects the learning and memory-related component of the underlying cognitive task. This overlap with general control functions is supported by the relatively strong correlation of factors 3 and 4.^31,75^

In summary, our results provide insight into common and distinct lesion-deficit patterns for language and executive control. The identified four factors of cognitive facets may inform future personalized cognitive therapy approaches optimized for individual lesion-deficit profile patterns. Individual deficit profile patterns may identify patients at risk for specific cognitive deficits early after stroke to enable targeted cognitive testing and therapy (^see 78^). While previous studies in patients with post-stroke aphasia suggested that recruitment of domain-general regions for cognitive control is linked to favorable language recovery^17–20^, none of these studies provided insight into the overlap of language and executive control functions. Our results fill this gap by showing that language-related control is associated with left-hemispheric regions while executive control is associated with the right hemisphere. Strikingly, the overlap of both functions is also mainly located in left-hemispheric regions. Yet, to date, individual lesion patterns are rarely considered for optimizing cognitive therapy. Based on our findings, we may speculate that lesion topologies associated with factor 2 may benefit most from language-specific treatment. In contrast, lesion topologies associated with factors 1 and 4 would additionally require the inclusion of more executive therapy elements.

## Abbreviations

BNT: Boston Naming Test
DSCT: Digit Symbol Coding Task
IQCODE: Informant Questionnaire on Cognitive Decline in the Elderly
MDN: multiple-demand network
MIMO: multiple-input multiple-output
MMSE: Mini Mental State Exam
MISO: multiple-input single-output
NMF: non-negative matrix factorization
RCF: Rey Complex Figure
SVL: Soul Verbal Learning
TMT: Trail Making Test

## Acknowledgments

The authors are grateful to Angelina K. Kancheva and Goezdem Arikan for their help with performing the manual infarct segmentations. DB was supported by the Brain Canada Foundation, through the Canada Brain Research Fund, with the financial support of Health Canada, National Institutes of Health (NIH R01 AG068563A, NIH R01 R01DA053301-01A1), the Canadian Institute of Health Research (CIHR 438531, CIHR 470425), the Healthy Brains Healthy Lives initiative (Canada First Research Excellence fund), Google (Research Award, Teaching Award), and by the CIFAR Artificial Intelligence Chairs program (Canada Institute for Advanced Research). GH was supported by Lise Meitner Excellence funding from the Max Planck Society, the European Research Council (ERC-2021-COG 101043747) and the German Research Foundation (HA 6314/3-1, HA 6314/4-2, HA 6314/9-1).

