## Supplementary Materials and Methods for "Bayesian modeling disentangles language versus executive control disruption in stroke"

**Participant recruitment**

We retrospectively selected 1401 patients from a stroke registry based on Bundang and Hallym Vascular Cognitive Impairment cohorts – prospectively recruited cohorts of patients initially admitted to the Seoul National University Bundang Hospital or Hallym University Sacred Heart Hospital in South Korea between 2007 and 2018^1^. These patients were diagnosed with acute ischaemic infarction based on diffusion-weighted MRI, typically within one week after symptom onset. Among all available patients, a total of 1080 patients were selected based on the following criteria: (1) availability of brain MRI showing acute tissue infarction in the diffusion-weighted imaging (DWI) and/or fluid-attenuated inversion recovery (FLAIR), (2) successful lesion segmentation and registration, (3) no previous cortical infarcts, large subcortical infarcts (>15 mm) or haemorrhages (>10 mm) on MRI, and (4) availability of follow-up data on key demographics and neuropsychological assessment (the 60-min Korean-Vascular Cognitive Impairment Harmonization Standards-Neuropsychology Protocol^2,3^. We excluded patients (1) whose MRI was inadequate for properly obtaining neuroimaging variables, (2) who had a bilateral stroke and (3) inability to undergo cognitive testing due to severe aphasia, as determined by the attending physician.

All subjects provided informed written consent in accordance with the Declaration of Helsinki. The local institutional review boards approved the study protocol and waived the required consent requirements based on the retrospective nature of this study and the minimal risk to participants. An identical participant sample was used in previously published research^4,5^.

**Neuroimaging data pre-processing**

Whole-brain MRI scans were typically acquired within the first week after the stroke event. Brain scanning included structural axial T1, T2-weighted spin echo, fluid-attenuated inversion recovery and DWI sequences (3.0 T, Achieva scanner, Philips Healthcare, Netherlands, image dimensions: 182 × 218 × 182; c.f., Neuroimaging protocols for details). Stroke lesions were manually segmented on DWI or, less frequently, FLAIR images by experienced, trained investigators (A.K.K. and G.A.) relying on in-house developed software based on MeVisLab (MeVis Medical Solutions AG, Bremen, Germany)^6^. Lesion segmentations were successively checked and potentially refined by two experienced raters (N.A.W. and J.M.B). Subsequently, images and corresponding lesion maps were linearly and non-linearly normalized to Montreal Neurological Institute (MNI-152) space employing the RegLSM image processing pipeline (public code: <http://lsm.isi.uu.nl/>)^7^. An experienced rater rigorously controlled the quality of normalization (N.A.W.). If there were any visual differences between the original and registered lesion maps during quality control, normalized lesion maps were manually corrected.

**Neuroimaging protocols**

(1) Seoul National University Bundang Hospital: The MRI protocols comprised diffusion-weighted imaging (DWI), axial T1- and T2weighted spin echo, fluid-attenuated inversion recovery imaging (FLAIR), gradient-echo imaging, and coronal T1-weighted spin echo imaging. FLAIR imaging was acquired using a fast-spin echo sequence with the acquisition parameters: repetition time: 11,000 ms; echo time: 125 ms; inversion time: 2800ms; slice thickness 5 mm; intersection gap 1mm; matrix: 512×512; flip angle 90 degrees. DWI imaging was obtained employing an EPI-spin echo sequence with the acquisition parameters: repetition time: 5000 ms; echo time: 50 ms; diffusion b-value: 1000; slice thickness: 5 mm; intersection gap: 1 mm; matrix: 256 × 256; flip angle 90 degrees. (2) Hallym University Sacred Heart Hospital: The MRI protocols comprised DWI, axial T1- and T2-weighted spin echo, FLAIR, gradient-echo imaging, and coronal T2-weighted spin echo imaging. FLAIR imaging acquisition parameters: repetition time: 11,000 ms; echo time: 125 ms; inversion time: 2800 ms; slice thickness: 5 mm; matrix: 512×512; flip angle 90 degrees. DWI image acquisition parameters: repetition time: 3000ms; echo time: 56ms; diffusion b-value: 1000; slice thickness: 5 mm; matrix: 256 ×256; flip angle 90 degrees.

**Atlas-based extraction of target lesion load signals**

Each subject was characterized by a total of 435 642 grey matter voxels of 1 mm^3^. To provide a more generalizable and interpretable form, we parsed each patient’s lesion fingerprint by summarizing the lesion load within 54 parcels (108 for both hemispheres) based on the Harvard-Oxford cortical atlas with 47 regions and subcortical atlas with 7 regions in each hemisphere^8^. We first counted the number of voxels affected per atlas-defined brain region. In doing so, we obtained 54 regional measures of lesion load per hemisphere in each participant. We then log-transformed and concatenated the ensuing lesion load measures for the left and right hemispheres.

**Data-driven deconvolution of lesion atoms**

After summarizing the lesion load across 54 anatomical brain regions for each subject, we sought to isolate coherent topographical patterns that may be hidden in these regional measures. To that end, we used non-negative matrix factorization (NMF)^9^ as a multivariate encoding strategy. NMF decomposed the input lesion map X of *n* subjects and *m* regions into two low-rank non-negative matrices W and H, such that:

$$X\approx WH,$$

where X is the [*m* x *n*] lesion load matrix, W is a [*m* x *k*] non-negative matrix for *k* latent patterns, and H is a non-negative matrix of size [*k* x *n*].

The matrix W represents a set of non-negative basis vectors (i.e., latent factor representations) that we will refer to as lesion atoms, which denotes region-wise implications in each of the *k* patterns. The latent pattern expression matrix H indicated how relevant each emerging lesion atom is to describe the constituent parts of an individual patient’s overall spatial lesion distribution in the brain. Specifically, our latent factorization deconvolved the actual lesion constellation in 10 unique combinations of spatially distributed region damage. The optimal number of 10 lesion atoms per hemisphere was validated in previous works on the identical dataset^5,10^. This global decomposition of local lesion load indicators strikes a balance between capturing a substantial amount of lesion variability, on the one hand, and keeping the number of quantities low for neuroscientific domain interpretation, on the other hand.

Embedding lesion load across the brain into 10 lesion atoms using NMF provided at least two key advantages to alternative dimensionality reduction tools. First, in contrast to clustering approaches that consider the effect of each location only once, each brain location could belong to several latent lesion components of W to varying degrees. In this way, each location could contribute to the prediction of cognitive scores through relative contributions of multiple components, each of which reflected extracted lesion archetypes distributed across the whole brain. As a second key advantage, the non-negativity of the segmented brain lesion information and the non-negativity constraint of the NMF model allowed for intuitive neurobiologically meaningful interpretations. That is, each latent component W_k_ represented a unique and directly interpretable aspect of the overall topographical lesion pattern variation. The neurobiologically interpretable sum-of-parts representation enabled by NMF constrasts with latent representations learned by alternative matrix factorization algorithms. For example, in principal component analysis, individual lesions would be recovered through convoluted additions and subtractions of several components with positive and negative weights. For this reason, the overall effect of all principal components, yet not the effect of ensuring individual components, would have been as easily and intuitively interpretable to draw neuroscientific conclusions.

**Latent factors driving cognitive outcomes**

Factor analysis re-represents the set of cognitive scores with a smaller set of hidden factors while maximizing the amount of explained correlation or, in other words, common variance (the amount of variance that is shared among cognitive scores). This common variance, along with unique variance (any portion of variance that is not shared among the 8 cognitive endpoints), makes up the total variance. In contrast, the commonly used principal components analysis assumes that there is no unique variance and the total variance is equal to the common variance.

The factor analysis model treats observed cognitive scores Y as measures of a smaller number of unobserved latent factors F, with corresponding loadings A, such that:

$$Y\approx FA'+E,$$

where Y is the [*n* × *l*] matrix of *l* centered (de-meaned across participants) and standardized (unit-variance scaled across participants) cognitive scores for *n* individuals, F is the [*n* × *p*] matrix of *n* individuals’ values for *p* latent variables, A is the [*l* × *p*] matrix of the latent variable effects on the *l* cognitive scores, and E represent score-specific [*n* x *l*] matrix of unique disturbance terms.

The weights A in the factor analysis express the relationship or association of each score Y to the underlying factor F. To estimate the parameters of the dimensionality reduction, the factor analysis finds a matrix of loadings A and a diagonal matrix Y such that the observed covariance matrix S is as well as possible approximated by:

$$\Sigma\approx AA'+\Psi$$

In other words, the factor analysis minimizes differences between off-diagonal elements of S and A. Since we can choose the diagonal matrix Y, the reconstruction error of the diagonal elements will be zero. While several methods exist for the extraction of latent factors, we selected the widely employed “varimax” solution that estimates factor loadings while minimizing the sum of squares of off-diagonal residuals^11^.

Due to rotational indeterminacy (an infinite number of equivalent A matrices up to a particular rotation), we chose the commonly used “varimax” rotation^12^. This method minimizes the number of variables that have high loadings on each factor and thus simplifies the interpretation of the factors. Based on the inspection of explained variance and associated eigenvalues, we opted for a four-factor solution. This solution reaches a compromise between the detailed and data-efficient characterization of the overarching cognitive dimensions. The derived underlying driving factors among our 8 target cognitive scores served as the basis for our modelling outcomes in downstream analyses.

**Bayesian Model specification**

Our generative, multi-level approach allowed us to obtain fully probabilistic parameter estimates that could inform us about effect strength and effect certainty of how each lesion pattern is responsible for the respective outcome. To directly examine possible differences in hemispheric predictive relevance for the selected outcome, the modelled generative process assumed a joint dispersion prior for all lesion atoms of each hemisphere. Therefore, the standard deviation priors for the left and right hemispheres could capture the hemisphere-specific predictive contributions tiled across all candidate lesion atoms. Priors of left- and right-hemispheric standard deviations were additionally combined through a joint hyperprior to complement the hierarchical model structure.

For all analytical solutions, samples from the joint posterior distribution of the model parameters were drawn by the No U-Turn Sampler, a Monte Carlo Markov Chain algorithm (setting: draws = 4000)^13^. Posterior predictive checks were carried out after model estimation to evaluate the obtained predictive model (with respect to its R^2^-based explained variance). In other words, we empirically assessed the simulated outcome predictions generated by our model solution to approximate external validation based on our patient sample. This empirical procedure is a well-recognized option for judging the adequacy of Bayesian models given the actual data at hand^14,15^.

**Full Bayesian model specification for MIMO model**

**Hyperpriors**

$$\boldsymbol{hype}\boldsymbol{r}_{\boldsymbol{\sigma}_{\boldsymbol{\beta}}}\boldsymbol{\sim Halfcauchy(\beta=1)}$$

$$\sigma_{\beta}\sim\boldsymbol{hype}\boldsymbol{r}_{\boldsymbol{\sigma}_{\boldsymbol{\beta}}}$$

**Priors**

$$\alpha\sim\boldsymbol{Normal(\mu=0, \sigma=1)}$$

$\beta_{1-10}\sim\boldsymbol{Normal(\mu=0, \sigma=}\boldsymbol{\sigma}_{\boldsymbol{\beta}}\boldsymbol{)}$

$\beta_{lesion load}\sim\boldsymbol{Normal(\mu=0, \sigma=10)}$

$\beta_{age}\sim\boldsymbol{Normal(\mu=0, \sigma=10)}$

$\beta_{{age}^{2}}\sim\boldsymbol{Normal(\mu=0, \sigma=10)}$

$\beta_{male}\sim\boldsymbol{Normal(\mu=0, \sigma=1)}$

$\beta_{female}\sim\boldsymbol{Normal(\mu=0, \sigma=1)}$

$\beta_{education years}\sim\boldsymbol{Normal(\mu=0, \sigma=5)}$

$\beta_{IQCODE}\sim\boldsymbol{Normal(\mu=0, \sigma=1)}$

$\beta_{time since onset}\sim\boldsymbol{Normal(\mu=0, \sigma=10)}$

**Likelihood of individual linear models (example for MMSE)**

$Y_{MMSE}=\alpha+\beta_{1-10}[hemisphere]+\beta_{lesion load}*Lesion load+\beta_{age}*age+\beta_{age^{2}}*age^{2}+$

$\beta_{male}*male+\beta_{female}*female+\beta_{education years}*education years+$

$\beta_{IQCODE}*IQCODE+\beta_{time onset}*time onse$t

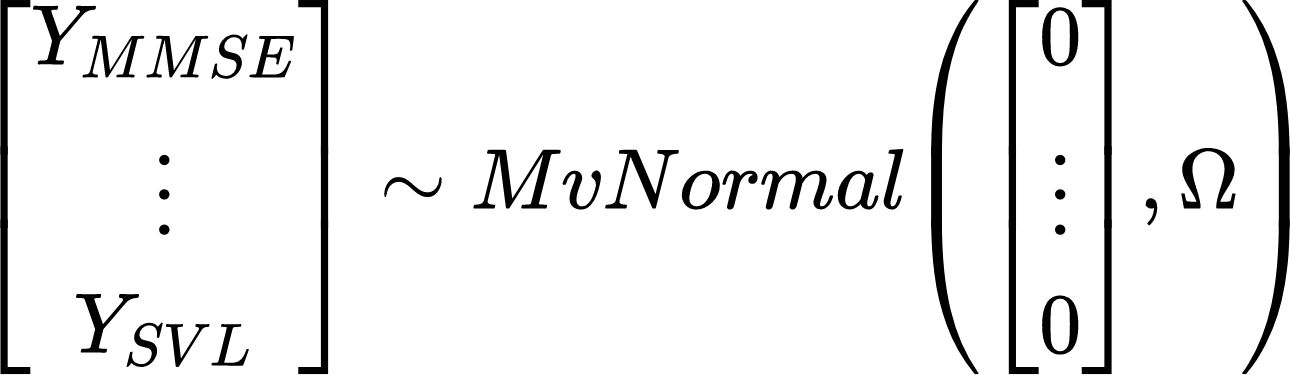

$$\Omega=LKJCorr(\eta=4, sd dist=Exponential(1))$$

**Full Bayesian model specification for MISO model (example for Factor 1)**

**Hyperpriors**

$$\boldsymbol{hype}\boldsymbol{r}_{\boldsymbol{\sigma}_{\boldsymbol{\beta}}}\boldsymbol{\sim Halfcauchy(\beta=1)}$$

$$\sigma_{\beta}\sim\boldsymbol{hype}\boldsymbol{r}_{\boldsymbol{\sigma}_{\boldsymbol{\beta}}}$$

**Priors**

$$\alpha\sim\boldsymbol{Normal(\mu=0, \sigma=1)}$$

$\beta_{1-10}\sim\boldsymbol{Normal(\mu=0, \sigma=}\boldsymbol{\sigma}_{\boldsymbol{\beta}}\boldsymbol{)}$

$\beta_{lesion load}\sim\boldsymbol{Normal(\mu=0, \sigma=10)}$

$\beta_{age}\sim\boldsymbol{Normal(\mu=0, \sigma=10)}$

$\beta_{{age}^{2}}\sim\boldsymbol{Normal(\mu=0, \sigma=10)}$

$\beta_{male}\sim\boldsymbol{Normal(\mu=0, \sigma=1)}$

$\beta_{female}\sim\boldsymbol{Normal(\mu=0, \sigma=1)}$

$\beta_{education years}\sim\boldsymbol{Normal(\mu=0, \sigma=5)}$

$\beta_{IQCODE}\sim\boldsymbol{Normal(\mu=0, \sigma=1)}$

$\beta_{time since onset}\sim\boldsymbol{Normal(\mu=0, \sigma=10)}$

**Likelihood of linear model**

$$Factor1=\alpha+\beta_{1-10}[hemisphere]+\beta_{lesion load}*Lesion load+\beta_{age}*age+\beta_{age^{2}}*age^{2}+$$

$\beta_{male}*male+\beta_{female}*female+\beta_{education years}*education years+$

$\beta_{IQCODE}*IQCODE+\beta_{time onset}*time onse$t

$$\varepsilon\sim Halfcauchy(\beta=20)$$

$$Factor1\sim Normal(\mu=Factor1,\sigma=\varepsilon)$$
